# HapCHAT: Adaptive haplotype assembly for efficiently leveraging high coverage in long reads

**DOI:** 10.1101/170225

**Authors:** Stefano Beretta, Murray D Patterson, Simone Zaccaria, Gianluca Della Vedova, Paola Bonizzoni

**Author notes:** Equal contributor (Stefano Beretta, Murray D Patterson).

## Abstract

**Background:** Haplotype assembly is the process of assigning the different alleles of the variants covered by mapped sequencing reads to the two haplotypes of the genome of a human individual. Long reads, which are nowadays cheaper to produce and more widely available than ever before, have been used to reduce the fragmentation of the assembled haplotypes since their ability to span several variants along the genome. These long reads are also characterized by a high error rate, an issue which may be mitigated, however, with larger sets of reads, when this error rate is uniform across genome positions. Unfortunately, current state-of-the-art dynamic programming approaches designed for long reads deal only with limited coverages.

**Results:** Here, we propose a new method for assembling haplotypes which combines and extends the features of previous approaches to deal with long reads and higher coverages. In particular, our algorithm is able to dynamically adapt the estimated number of errors at each variant site, while minimizing the total number of error corrections necessary for finding a feasible solution. This allows our method to significantly reduce the required computational resources, allowing to consider datasets composed of higher coverages. The algorithm has been implemented in a freely available tool, HapCHAT: <u>Hap</u>lotype Assembly <u>C</u>overage <u>H</u>andling by <u>A</u>dapting <u>T</u>hresholds. An experimental analysis on sequencing reads with up to 60× coverage reveals improvements in accuracy and recall achieved by considering a higher coverage with lower runtimes.

**Conclusions:** Our method leverages the long-range information of sequencing reads that allows to obtain assembled haplotypes fragmented in a lower number of unphased haplotype blocks. At the same time, our method is also able to deal with higher coverages to better correct the errors in the original reads and to obtain more accurate haplotypes as a result.

**Availability:** HapCHAT is available at http://hapchat.algolab.eu under the GPL license.

## Introduction

Due to the diploid nature of the human genome, *i.e*., it has two copies of its genome, called *haplotypes*, genomic variants appear on either of these two copies. Knowing the specific haplotype on which each of the genomic variants occurs has a strong impact on various studies in genetics, from population genomics [1, 2], to clinical and medical genetics [3], or to the effects of compound heterozygosity [2, 4].

More specifically, the variations between two haplotypes of the genome are, for the most part, in the form of heterozygous Single Nucleotide Variants (SNVs), i.e., single genomic positions where the haplotypes contain two distinct alleles. Since a direct experimental reconstruction of the haplotypes is not yet cost effective [5] or require methods that have not yet gained widespread adoption [6, 7], computational methods aim to perform this task starting from sequencing reads mapped to a reference human genome. In fact, sequencing reads usually cover multiple SNV positions on the genome, hence providing information about the corresponding alleles that co-occur on a haplotype. In particular, *haplotype assembly* is the computational approach aiming to partition the reads into two sets such that all the reads belonging to the same set are assigned to the same haplotype.

Due to the availability of curated, high quality haplotype reference panels on a large population of individuals [8, 9], computational methods for statistically inferring the haplotypes of an individual from these panels are widely used [10, 1]. The accuracy of these methods, however, depends heavily on the size and diversity of the population used to compile the panels, entailing poor performance on rare variants, while *de novo* variants are completely missed. These types of variants appear in the sequencing reads of the individual, making read-based haplotype assembly the obvious solution.

The combinatorial *Minimum Error Correction* (MEC) problem is the most commonly cited formulation of haplotype assembly [11]. Under the principle of parsimony, MEC aims to find the minimum number of corrections to the values of sequencing reads in order to be able to partition the reads into two haplotypes. Unfortunately, this problem is NP-hard [11] and it is even hard to approximate [12, 13, 14]. As such, several heuristics for haplotype assembly have been proposed [15, 16, 17, 18, 19]. Beyond that, several exact methods have been proposed, including *Integer Linear Programming* (ILP) approaches [20, 21], and *Dynamic Programming* (DP) approaches which are *Fixed-Parameter Tractable* (FPT) in some parameter [22, 13]. These methods achieve good results on datasets obtained using the traditional short sequencing reads. However, short reads do not allow to span more than a few SNV positions along the genome, rendering them inadequate for reconstructing long regions of the two haplotypes. In fact, the short range information provided by these reads does not allow to link many – if any – SNVs together. Consequently, the resulting assembled haplotypes are fragmented into many short haplotype blocks that remain unphased, relative to each other [23].

The advent of third generation sequencing technologies introduces a new kind of sequencing reads, called *long reads*, that are able to cover much longer portions of the genome [24, 25, 26]. Each read may span several positions along the genome and the long-range information provided by these reads allow to link several SNVs. This results in the possibility of obtaining longer haplotype blocks that assign more variants to the corresponding haplotype [27, 28]. Current third generation sequencing platforms offered by Pacific Biosciences (PacBio) [29] and Oxford Nanopore Technologies (ONT) [30] are now able to produce reads of tens to hundreds of kilo-basepairs (kbp) in length, and are much more capable of capturing together more variants than the short reads that are commonplace today. While PacBio technologies are characterized by a high error rate (substitution error rate up to 5% and indel rate up to 10%), this is uniformly distributed along the genome positions [24, 25, 31] – something we can take advantage of. Oxford Nanopore Technologies, on the other hand, have an even higher error rate which is also not uniformly distributed [32]. Traditional approaches that have been designed for short reads fail when they are applied to these long reads, even when considering low coverages, as demonstrated in [33]. This is due to the fact that these approaches scale poorly with increasing read length [21, 22].

Recently, two methods have been proposed to specifically deal with long reads and their characteristics, namely WhatsHap [33, 34] and HapCol [35]. On the one hand, WhatsHap introduces a dynamic programming algorithm that is fixed parameter tractable, with *coverage* as the parameter, where coverage is the maximum number of reads covering any genome position. Hence, this algorithm is able to leverage the long-range information of long reads since its runtime is independent of the read length, but unfortunately it can deal only with datasets of limited coverages – up to 20×, and hence resorts to pruning datasets with higher coverage [33]. A parallel version of WhatsHap has been recently proposed showing the capability to deal with higher coverages of up to 25× [36]. Although WhatsHap computes the theoretically optimal solution to the MEC problem, minimizing the overall number of corrections in the input reads, this could result, however, in columns having an unrealistically large number of corrections, which may not be coherent with how the errors are truly distributed in the actual reads.

On the other hand, HapCol proposes an approach that exploits the uniform distribution of sequencing errors characterizing long reads. In particular, the authors propose a new formulation of the MEC problem where the maximum number of corrections is bounded in every column and is computed from the expected error rate [35]. HapCol has been shown to be able to deal with datasets of higher coverages compared to WhatsHap. However, the presence of genome positions containing more errors than expected (due to errors in the alignment or repetitive regions) is a problem for this approach. As a result, even HapCol was effectively limited to deal with instances of relatively low coverages up to 25–30×, since even the presence of few outliers forces the algorithm to change the global behavior, or to fail.

As a result, both the methods proposed for haplotype assembly from long reads, WhatsHap and HapCol, have issues managing datasets with increasing coverages. However, considering a higher number of reads covering each position is indeed the most reliable way to face the high error rate characterizing the sequencing reads produced by third generation sequencing technologies. In fact, long reads generated by the PacBio platform share a limited number of errors on any given SNV position that they cover because errors are almost uniformly distributed across genome positions. Therefore, increasing the coverage mitigates the effects of sequencing errors and may allow to reconstruct haplotypes of higher quality.

In this work we propose a new method which combines and extends the main features of the previous WhatsHap and HapCol, and aims to deal with datasets of higher coverages while being robust to the presence of noise and outliers. In particular, we re-design the approach proposed in [35] by allowing also the dynamic adaption of the estimated error rate and, consequently, the maximum number of corrections that are allowed in each position. This allows the handling of columns that require more errors than expected, while avoiding the exploration of scenarios that involve a number of corrections that is much higher than necessary for a site. This is coupled with a merging procedure which merges pairs of reads that are highly likely to originate from the same haplotype, allowing this method to scale to significantly higher values of coverage. The method has been implemented in HapCHAT: <u>Hap</u>lotype Assembly <u>C</u>overage <u>H</u>andling by <u>A</u>dapting <u>T</u>hresholds that is freely available at http://github.com/AlgoLab/HapCHAT. An experimental analysis on real and simulated sequencing reads with up to 60× coverage reveals that we are able to leverage high coverage towards better predictions in terms of both accuracy (switch error rate) and recall (QAN50 score), as we see an upward trend in both, as coverage increases. This trend is the most stark in the case of recall, which is where it counts the most, since the ultimate goal of haplotype assembly is indeed to assemble the longest haplotype blocks possible.

We compare our method to some of the state-of-the-art methods in haplotype assembly, including HapCol [35]; the newest version of WhatsHap [37], to which many features have since been added; and HapCUT2 [16, 17]. We show that HapCHAT is comparable to or better than any tool in terms of both accuracy and recall, while requiring an amount of computational resources (time and memory) that is on the same or a lower order of magnitude of any comparable (in terms of accuracy or recall) tool in every case. These results confirm that high coverage can indeed be leveraged in order to deal with the high error rate of long reads in order to take advantage of their long-range information.

## Background

Let *υ* be a vector, then *υ*[*i*] denotes the value of *υ* at position *i*. A *haplotype* is a vector *h* ∈ {0,1}^*m*^. Given two haplotypes of an individual, say *h*_1_, *h*_2_, the position *j* is *heterozygous* if *h*_1_[*j*] ≠ *h*_2_[*j*], otherwise *j* is *homozygous*. A *fragment* is a vector *f* of length *l* belonging to {0,1, −}^*l*^. Given a fragment *f*, position *j* is a *hole* if *f*[*j*] = −, while a *gap* is a maximal sub-vector of *f* of holes, *i.e*., a gap is preceded and followed by a non-hole element (or by a boundary of the fragment).

A *fragment matrix* is a matrix *M* that consists of *n* rows (fragments) and *m* columns (SNVs). We denote as *L* the maximum length for all the fragments in *M*, and as *M_j_* the *j*-th column of *M*. Notice that each column of *M* is a vector in {0,1, −}^*n*^ while each row is a vector in {0,1, −}^*m*^.

Given two row vectors *r*_1_ and *r*_2_ belonging to {0,1, −}^*m*^, *r*_1_ and *r*_2_ are in *conflict* if there exists a position *j*, with 1 ≤ *j* ≤ *m*, such that *r*_1_[*j*] ≠ *r*_2_[*j*] and *r*_1_[*j*], *r*_2_[*j*] ≠ −, otherwise *r*_1_ and *r*_2_ are in *agreement*. A fragment matrix *M* is *conflict free* if and only if there exist two haplotypes *h*_1_, *h*_2_ such that each row of *M* is in agreement with one of *h*_1_ and *h*_2_. Equivalently, *M* is conflict free if and only if there exists a *bipartition* (*P*_1_, *P*_2_) of the fragments in *M* such that each pair of fragments in *P*_1_ is in agreement and each pair of fragments in *P*_2_ is in agreement. A *k-correction* of a column *M_j_*, is obtained from *M_j_* by flipping at most *k* values that are different from –. A column of a matrix is called *homozygous* if it contains no 0 or no 1, otherwise (if it contains both 0 and 1) it is called *heterozygous*. We say that a fragment *i* is *active* on a column *M_j_*, if *M_j_*[*i*] = 0 or *M_j_*[*i*] = 1. The *active fragments* of a column *M_j_* are the set *active* (*M_j_*) = {*i*: *M_j_*[*i*] ≠ −}. The *coverage* of the column *M_j_* is defined as the number *cov_j_* of fragments that are active on *M_j_*, that is *cov_j_* = |*active*(*M_j_*)|. In the following, we indicate as *cov* the maximum coverage over all the columns of M. Given two columns *M_i_* and *M_j_*, we denote by *active* (*M_i_, M_j_*) the intersection *active* (*M_i_*) ∩ *active* (*M_j_*). Moreover, we will write *M_i_* ≈ *M_j_*, and say that *M_i_, M_j_* are in accordance [13], if *M_i_*[*r*] = *M_y_*[*r*] for each *r* ∈ *active*(*M_i_, M_j_*), or *M_i_*[*r*] ≠ *M_y_*[*r*] for each *r* ∈ *active*(*M_i_, M_j_*). Notice that *M_i_* ≈ *M_j_* means that these two columns are compatible, that is, they induce no conflict. Moreover, *d*(*M_i_, M_j_*) denotes the minimum number of corrections to make columns *M_i_* and *M_j_* in accordance.

The *Minimum Error Correction* (MEC) problem [11, 38], given a matrix *M* of fragments, asks to find a conflict free matrix *C* obtained from *M* with the minimum number of corrections. In this work, we consider the variant of the MEC problem, called *k*-cMEC in which the number of corrections per column is bounded by an integer *k* [35]. More precisely, we want a *k-correction matrix D* for *M* where each column *C_j_* is a *k*-correction of column *M_j_*, minimizing the total number of corrections. We recall that in this paper we will consider only matrices where all columns are heterozygous.

Now, let us briefly recall the dynamic programming approach to solve the *k*-cMEC problem [35]. This approach computes a bidimensional array *D*[*j, C_j_*] for each column *j* ≥ 1 and each possible heterozygous *k*-correction *C_j_* of *M_j_*, where each entry *D*[*j, C_j_*] contains the minimum number of corrections to obtain a *k*-correction matrix *C* for *M* on columns *M*_1_,…, *M_j_* such that the columns *C_j_* are heterozygous. For the sake of simplicity, we pose *D*[0, ·] = 0. For 0 < *j* ≤ *m*, the recurrence equation for *D*[*j,C_j_*] is the following, where *δ_j_* is the set of all heterozygous *k*-corrections of the column *M_j_*.

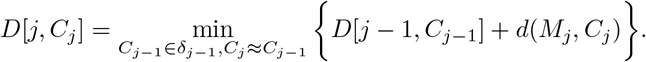

For the complete description of the dynamic programming recurrence we refer the reader to [35, 13].

## Methods

In this section, we highlight the new insight of HapCHAT for the assembly of single individual haplotypes, with the specific goal of processing high coverage in long read datasets. In fact, as reported in the original HapCol paper [35], the FPT algorithm is exponential in the number *k* of allowed corrections in each position. Therefore, we developed a preprocessing step which merges reads belonging to the same haplotype based on a graph clustering method. Moreover, we also improved the HapCol method by introducing a heuristic procedure to cope with problematic positions, i.e. those requiring more than *k* corrections.

As anticipated, the combination of all these improvements allowed the possibility of reconstructing haplotypes using higher coverage reads (w.r.t. the original HapCol method), while reducing the runtimes.

### Preprocessing

The first step of our pipeline is to merge pairs of fragments that, with high probability, originate from the same haplotype. With *p* we denote the (average) probability that any single base has been read incorrectly (i.e. that a nucleotide in the input BAM file is wrong) — we recall that *p* ≈ 0.15 and that errors are uniformly distributed for PacBio reads. Let *r*_1_ and *r*_2_ be two reads that share *m* + *x* sites, where they agree on *m* of those sites and disagree on the other x sites. For this pair of reads, we compute a likelihood under the hypothesis that the reads originate from the same haplotype, and a likelihood under the hypothesis that the reads originate from different haplotypes. We then compute the ratio of these two likelihoods. This idea is similar to the one adopted in [39], but our use is different.

Then, the probability of obtaining the two reads *r*_1_ and *r*_2_ under the hypothesis that they originate from the same haplotype is approximately *p_s_*(*r*_1_,*r*_2_) = (1 – *p*)^2*m*^*p^x^*(1 – *p*/3)^*x*^, that is we assume that we have no error in the shared part and exactly one error on the other sites. Similarly, the probability of obtaining the two reads *r*_1_ and *r*_2_ under the hypothesis that they originate from two different haplotypes is approximately *p_d_*(*r*_1_,*r*_2_) = *p^m^*(1 – *p*/3)^*m*^(1 – *p*/3)^*x*^(1 – *p*)^*x*^, that is we assume that there is exactly one error in the sites with same value and at most an error in the sites with different values.

A simple approach to reduce the size of the instance is to merge all pairs (*r*_1_, *r*_2_) of fragments such that *p_s_*(*r*_1_,*r*_2_) is sufficiently large. But that would also merge some pairs of fragments whose probability *p_d_* is too large. Since we want to be conservative in merging fragments, we partition the fragment set into clusters such that *p_s_/p_d_* ≥ 10^6^ for each pair of fragments in the cluster. This threshold was obtained empirically, in order to achieve the best performance in terms of quality of the predictions in the performed experimental analysis. Then, for each site, the character that is the result of a merge is chosen applying a majority rule, weighted by the Phred score of each symbol. Notice that the merging heuristic of ProbHap [39] considers only the ratio to determine when to merge two reads, while we analyze all pairs of reads to determine which *sets* of reads to merge.

### Adaptive k-cMEC

Here, we describe how we modified the HapCol dynamic programming recurrence in order to deal with problematic columns for which the maximum allowed number of corrections is not enough to obtain a solution. As stated in the original HapCol paper [35], the number *k_j_* of corrections for each column *M_j_* is computed, based on its coverage *cov_j_* and on two input parameters: *∊* (average *error-rate*) and *α* (the probability that the column *M_j_* has more than *k_j_* errors). The idea is that the number of errors in a column *j* follows a binomial distribution, and hence we allow the lowest value *k_j_* such that the probability of having more than *k_j_* errors (with error rate *∊*) is at most *α*. This is done in order to bound the value of *k*, which is fundamental since HapCol implements an FPT algorithm that is exponential in the maximum number of allowed corrections. For this reason, we would prefer to have low values of *k_j_*. A side effect of this approach is that, when all solutions of an instance contain a column with more than *k_j_* errors, HapCol is not able to find a solution. Therefore, we developed a heuristic procedure which has the final goal of guaranteeing that a solution is found, by slightly increasing the allowed number of errors beyond *k_j_*, such that a solution exists for this number. We recall that the recurrence equation governing the original dynamic programming approach considers all *k_j_*-correction *C_j_* ∈ *δ_j_*. We slightly modify the definition of *k*-corrections to cope with those problematic columns, by increasing the number of allowed corrections. Let *C_j,k_* be a *k*-correction of *M_j_* with exactly *k* corrections, let *z*_*j*,0_ = *k_j_* and *z_j,i_* = *z*_*j*,*i*−1_ + ⌊log_2_(*z_j,i-1_*) + 1⌋, *i.e*., each term is obtained from the previous one by adding a logarithmic term, to guarantee that the number of allowed corrections does not grow too quickly. Then 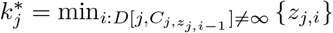 if *i* > 0, where *D*[·, ·] ≠ ∞ means it is a feasible correction. Starting from this notation, the new set of possible corrections of column *M_j_* is

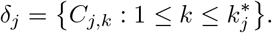

Notice that the sequence of *z_j,i_* is monotonically increasing with *i*, hence we can compute 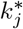 by starting with *k_j_* and increasing it until we are able to find a 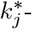 – correction for the column *M_j_*. The dynamic programming equation is unchanged, but our new construction of the set *δ_j_* guarantees that we are always able to compute a solution. Moreover, just as for HapCol, we cannot guarantee that we solve optimally the instance of the MEC problem.

One of the key points of this procedure is how we increment *z_j,i_*, that is by adding a logarithmic quantity. This guarantees a balance between finding a low value of 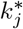 and the running time needed for the computation.

## Results and Discussion

We now describe the results of our experiments. In the first subsection, we describe the data that we use, or simulate. Then we detail the experiments that we set up in order to compare our tool with others in the next subsection. Finally, we present and discuss the results of these experiments.

### Data description

The Genome in a Bottle (GIAB) Consortium has released publicly available high-quality sequencing data for seven individuals, using eleven different technologies [40, 41, 42]. Since our goal is to assess the performance of different single-individual haplotype phasing methods, we study chromosome 1 of the Ashkenazim individual NA24385, as well as chromosomes 1-22 of individual NA12878.

The Ashkenazim individual is the son in a mother-father-son trio. We downloaded from GIAB the genotype variants call sets NIST_CallsIn2Technologies_05182015, a set of variants for each individual of this trio that have been called by at least two independent variant calling technologies. In order to be able to compare against methods that use reference panels or information from multiple individuals, *e.g*., a trio, for single-individual haplotype phasing, we considered all the bi-allelic SNVs of the chromosome that: (a) appear also in the 1000 Genomes reference panel https://mathgen.stats.ox.ac.uk/impute/1000GP_Phase3.tgz, and (b) have been called in all three individuals of the Ashkenazim trio, *i.e*., also in the mother and the father. For chromosome 1, this resulted in 140744 SNVs, of which 48023 are heterozygous. We refer to this set of SNVs as the set of benchmark SNVs for this dataset – the set is in the form of a VCF file. Since the authors of [43] also studied this trio, and have made the pipeline for collecting and generating their data publicly available at https://bitbucket.org/whatshap/phasing-comparison-experiments/, we use or modify parts of this pipeline to generate our data as detailed in the following.

As for the individual NA12878, we downloaded the latest high confidence phased VCF of GIAB for hg37, available at ftp://ftp-trace.ncbi.nlm.nih.gov/giab/ftp/release/NA12878_HG001/latest/GRCh37/HG001_GRCh37_GIAB_highconf_CG-IllFB-IllGATKHC-Ion-10X-SOLID_CHROM1-X_v.3.3.2_highconf_PGandRTGphasetransfer.vcf.gz, and used all SNVs in this file as our set of benchmark SNVs for the respective chromosomes.

### GIAB PacBio Reads

One of the more recent technologies producing long reads – those which are the most informative for read-based phasing – is the Pacific Biosciences (PacBio) platform. PacBio is one of the eleven technologies on which GIAB provides sequencing reads.

We hence downloaded the set of aligned PacBio reads from ftp://ftp-trace.ncbi.nlm.nih.gov/giab/ftp/data/AshkenazimTrio/HG002_NA24385_son/PacBio_MtSinai_NIST/MtSinai_blasr_bam_GRCh37/hg002_gr37_1.bam for chromosome 1 of the Ashkenazim individual, which has an average coverage of 60.2× and an average mapped read length of 8687 bp. We then downsampled the read set to average coverages of 25×, 30×, 35×, 40×, 45×, 50×, 55×, and 60×. This was done using the DownsampleSam subcommand of Picard Tools, which randomly downsamples a read set by selecting each read with probability *p*. We downsample recursively, so that each downsampled read set with a given average coverage is a subset of any downsampled read set with an average coverage higher than this set.

As for individual NA12878, we downloaded the set of aligned PacBio reads ftp://ftp-trace.ncbi.nlm.nih.gov/giab/ftp/data/NA12878/NA12878_PacBio_MtSinai/sorted_final_merged.bam, which comprises chromosomes 1–22. The average coverages (resp., mapped read lengths) ranged between 26.9 and 44.2 (resp., 4746 and 5285), so we did not perform any downsampling for this dataset.

As a phasing benchmark for the Ashkenazim chromosome 1, we used the latest high confidence trio-phased VCF of GIAB for hg37, available at ftp://ftp-trace.ncbi.nlm.nih.gov/giab/ftp/release/AshkenazimTrio/HG002_NA24385_son/latest/GRCh37/HG002_GRCh37_GIAB_highconf_CG-IllFB-IllGATKHC-Ion-10X-SOLID_CHROM1-22_v.3.3.2_highconf_triophased.vcf.gz. As for chromosomes 1–22 of the individual NA12878, we used the (original, *i.e*., phased version of the) high confidence phased VCF mentioned in the previous section.

### Simulated PacBio Data

Aside from the PacBio data described in the previous section, we also produce and run our experiments on a simulated read set for chromosome 1 of the Ashkenazim individual. Reference panels may leave out some variants with low allele frequency – a good reason for doing read-based phasing – and statistical methods might be susceptible to systematic bias in the data. For these reasons, we complement our study with an experimental analysis on simulated reads, as follows.

We first obtain a pair of “true” haplotypes off of which we simulate reads. This is obtained from the output of the population-based phasing tool SHAPEITv2-r837 [44] with default parameters on the 1000 Genomes reference panel, the corresponding genetic map http://www.shapeit.fr/files/genetic_map_b37.tar.gz, and the unphased genotypes, *i.e*., the set of benchmark SNVs of this chromosome.

Given the phasing by SHAPEIT, we incorporate the (benchmark) SNVs of the first haplotype of this phasing into the reference genome (hg37) by flipping the variant sites that are the alternative allele in this haplotype. The second haplotype is obtained analogously. Using these two true haplotypes as the input, we produce a corresponding set of reads for this haplotype using PBSIM [45], a PacBio-specific read simulator. We input to PBSIM the optional parameters `--depth` 60 so that our simulated reads have sufficient coverage, and as `--sample-fastq` a sample of the original GIAB PacBio reads described in the previous section, so that our simulated reads have the same length and accuracy profile as the corresponding real read set. We align the resulting simulated reads to the reference genome using BWA-MEM 0.7.12-r1039 [46] with optional parameter `–x pacbio`. Finally, this pair of aligned read sets, representing the reads coming off of each haplotype is merged using the `MergeSamFiles` subcommand of Picard Tools, obtaining the final simulated read set. In the same way as we have done with the read sets for the real Chromosome 1, we downsample to average coverages 25×, 30×, 35×, 40×, 45×, 50×, 55×, and 60×.

To summarize, the data we use or simulate regards both real and simulated reads on chromosome 1 of the Ashkenazim individual for a set of 8 average coverages, for a total of 16 read sets, each in the form of a BAM file. The autosomes of individual NA12878 adds an additional 22 read sets, each in the form of a BAM file. It is on these 38 read sets, along with their corresponding set of benchmark SNVs – in the form of VCF files – that we carry out our experiments, as described in the following section.

### Experimental Setup

We compare our tool HapCHAT to the most recent state-of-the-art read-based phasing methods of WhatsHap [34, 37], HapCol [35], HapCUT2 [17], ProbHap [39], ReFHap [19] and FastHare [15] by running them all on the data described in the previous subsection. Recall that, as detailed in the introduction, WhatsHap, HapCol and HapCHAT are approaches with a core phasing algorithm that is FPT either in the coverage or in the number of errors at each SNV site. Hence the coverage must first be reduced to some target maximum coverage before its core algorithm can be run. Each run of a tool on a dataset is given a time limit of one day, and a memory limit of 64GB. We now describe the details of how we parameterized each tool for comparison in what follows.

### WhatsHap

For each read set, we provide to WhatsHap (version 0.13) the corresponding BAM and VCF file. We run WhatsHap on this input pair on otherwise default settings, with the exception of providing it the reference genome (hg37) via the optional parameter `--reference`. This allows WhatsHap to run in *realignment mode*, which has been shown to significantly boost accuracy predictions for noisy read sets such as PacBio, as detailed in [37]. In particular, this mode is well suited to handle the abundant indel errors in the input reads. WhatsHap has a built-in read selection procedure [47] which subsequently prunes to a default maximum coverage of 15 before the core phasing algorithm is called. The default value has been selected by the authors of WhatsHap to provide the best trade-off between quality of the results and runtime [48]. Additionally, we run WhatsHap in realignment mode as above, but fixing to target maximum coverage 20 by providing the additional optional parameter `–H 20`. It is the resulting set of phasings by WhatsHap, in the form of phased VCF, that we use for the basis of comparison with the other methods.

### HapCol

For each read set, together with the VCF file of the corresponding chromosome, we convert it to the custom input format for HapCol. Since HapCol does not have a read selection procedure – something it does need for data at 35× (or higher) coverage (*cf*. the Introduction) — we then apply the read selection procedure of [47] to prune this set to the target maximum coverages of 15×, 20×, 25×, and 30×. On these resulting input files, we run HapCol with its default value of *α* = 0.01 (and of *ε* = 0.05) (*cf*. the subsection on Adaptive *k*-cMEC or [35] for details on the meaning of *α* and *ε*). Since HapCol is not adaptive, but we want to give it a chance to obtain a solution on its instance, should a given *α* be infeasible (*cf*. the subsection on Adaptive *k*-cMEC), we continue to rerun HapCol with an *α* of one tenth the size of the previous until a solution exists. HapCol outputs a pair of binary strings representing the phasing, which we then convert to phased VCF. Note that we did not further attempt any higher maximum coverages, because at maximum coverage 30, HapCol either exceeded one day of runtime or 64GB of memory on every dataset. It is this set of resulting phasings (phased VCF files) that we use to compare with the other methods.

### ProbHap, RefHap and FastHare

For each read set, we use the `extractHAIRS` program that is distributed with the original HapCut [16] to convert its BAM / VCF pair into the custom input format for these methods. We then ran each method on these instances with default settings, each producing a custom input which is then converted to a phased VCF with the subcommand `hapcut2vcf` of the WhatsHap toolbox.

### HapCUT2

For each read set, we use the `extractHAIRS` program that comes with HapCUT2, with parameter `--pacbio 1`, which activates a newly-developed realignment procedure for pacbio reads, to convert its BAM / VCF pair into the custom input format for HapCUT2. We then ran HapCUT2 on the resulting instances with default settings, each producing a custom output which is then converted to phased VCF with the subcommand `hapcut2vcf` of the WhatsHap toolbox.

### HapCHAT

For each read set, we provide to HapCHAT the corresponding BAM and VCF file. We run HapCHAT on this input pair on otherwise default settings, with the exception of providing it the reference genome (hg37) via the optional parameter `--reference`. This allows HapCHAT to run in realignment mode like with WhatsHap, thanks to the partial integration of HapCHAT into the WhatsHap codebase. We then apply our merging step as described in the subsection Preprocessing, which reduces the coverage. If necessary, the reads are further selected via a greedy selection approach (based on the Phred score), with ties broken at random, to downsample each dataset to the target maximum coverages of 15×, 20×, 25×, and 30×. It is the resulting phasings, in phased VCF format, for which the comparison of HapCHAT to other methods is based.

### Experimental results and discussion

The times reported here do not include the time necessary to read the input (BAM) file, which is more-or-less the same for each method. The results are summarized in Tables 1–17 and Figures 1,2 and 3.

**Figure 1.**
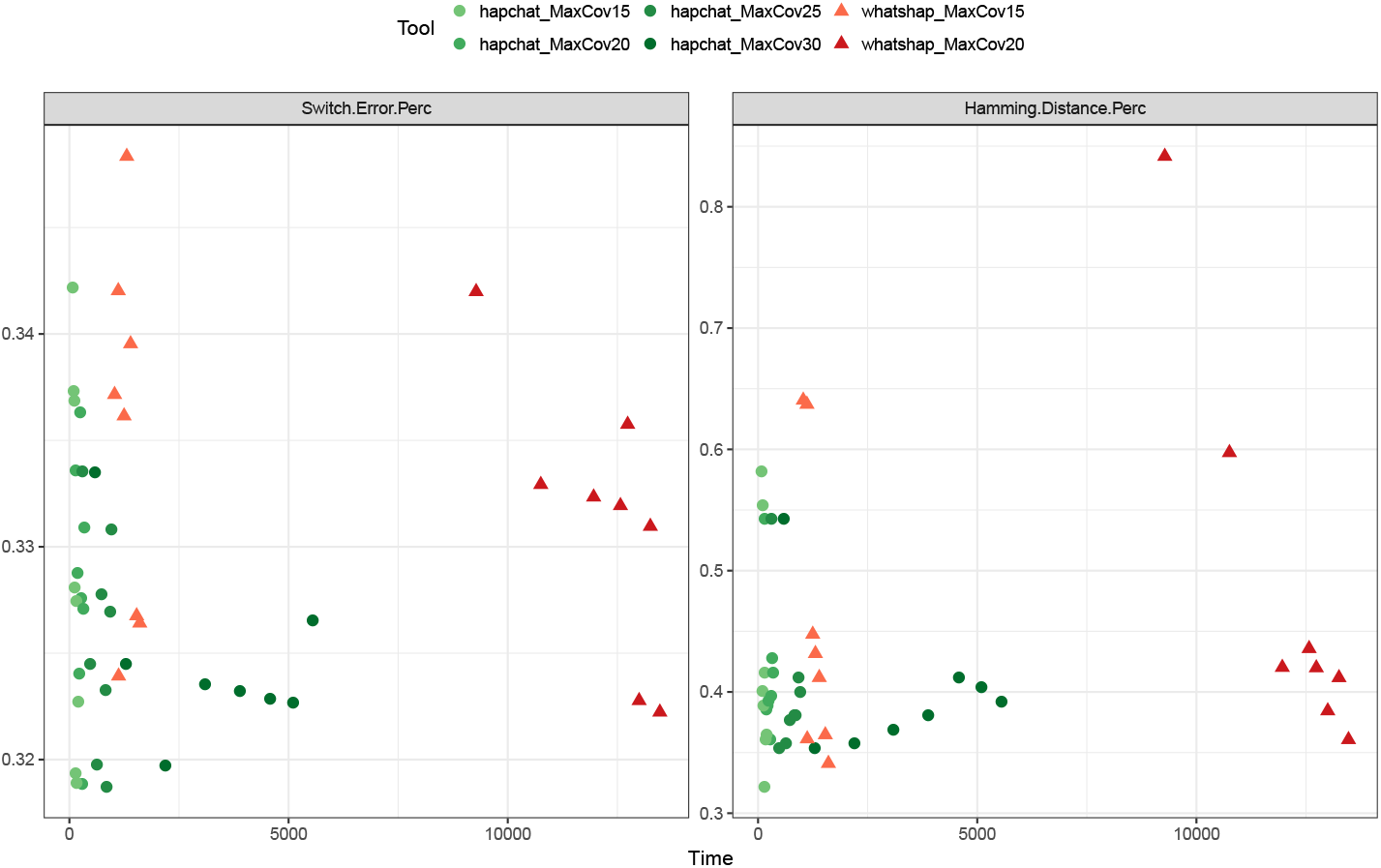
Switch error rate and Hamming distance as a function of running time. As achieved by HapCHAT and WhatsHap at different maximum coverages on the real Ashkenazim Chromosome 1 dataset. For each tool and each maximum coverage, we represent a point for each of the 8 possible values of the average coverage.

**Figure 2.**
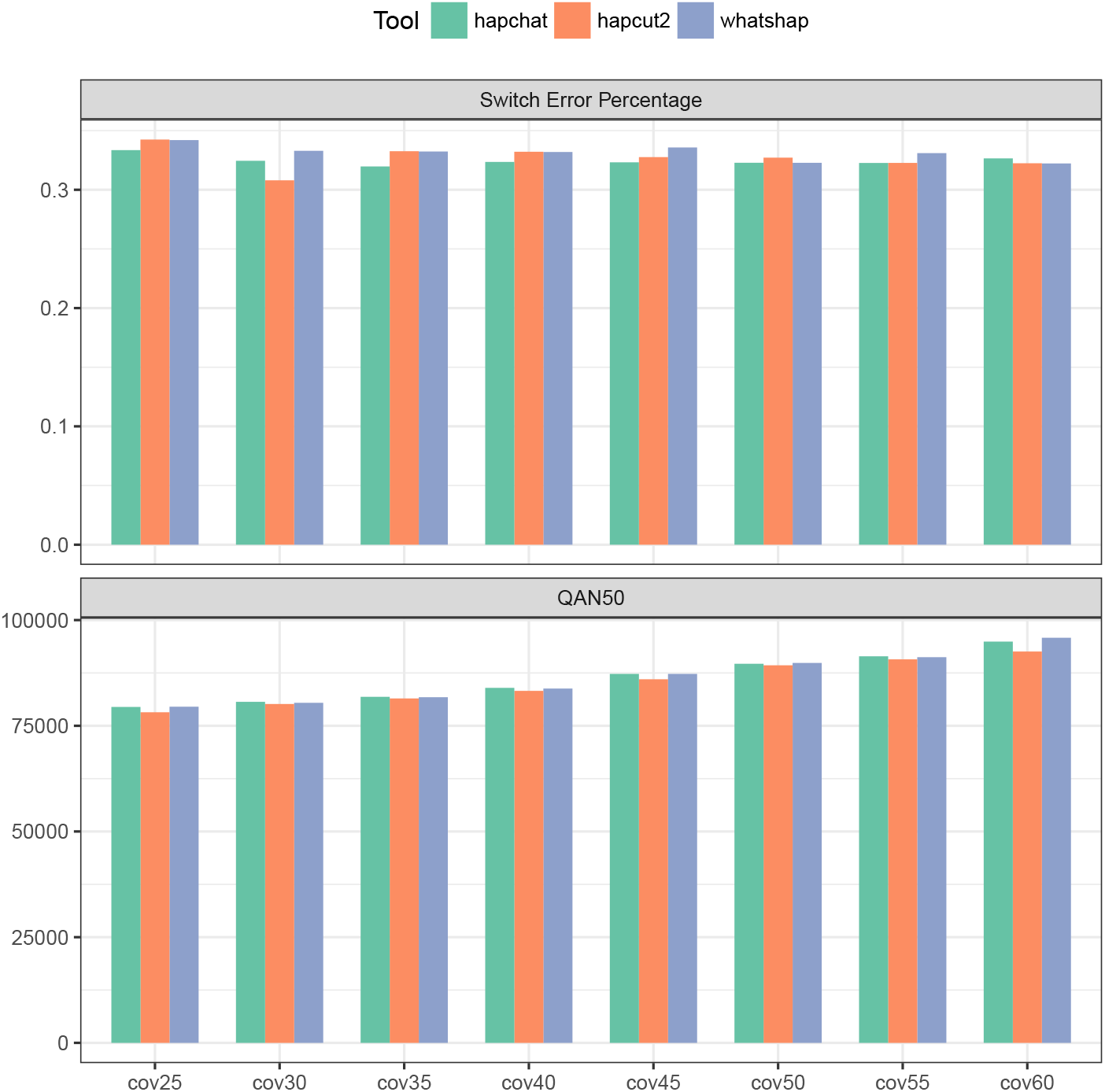
Quality measures on the real Ashkenazim Chromosome 1 dataset. We present the bar plots showing the measures of switch error percentage and QAN50 achieved by HapCHAT, WhatsHap, and HapCUT2 on the Ashkenazim Chromosome 1 dataset at different coverage values.

**Figure 3.**
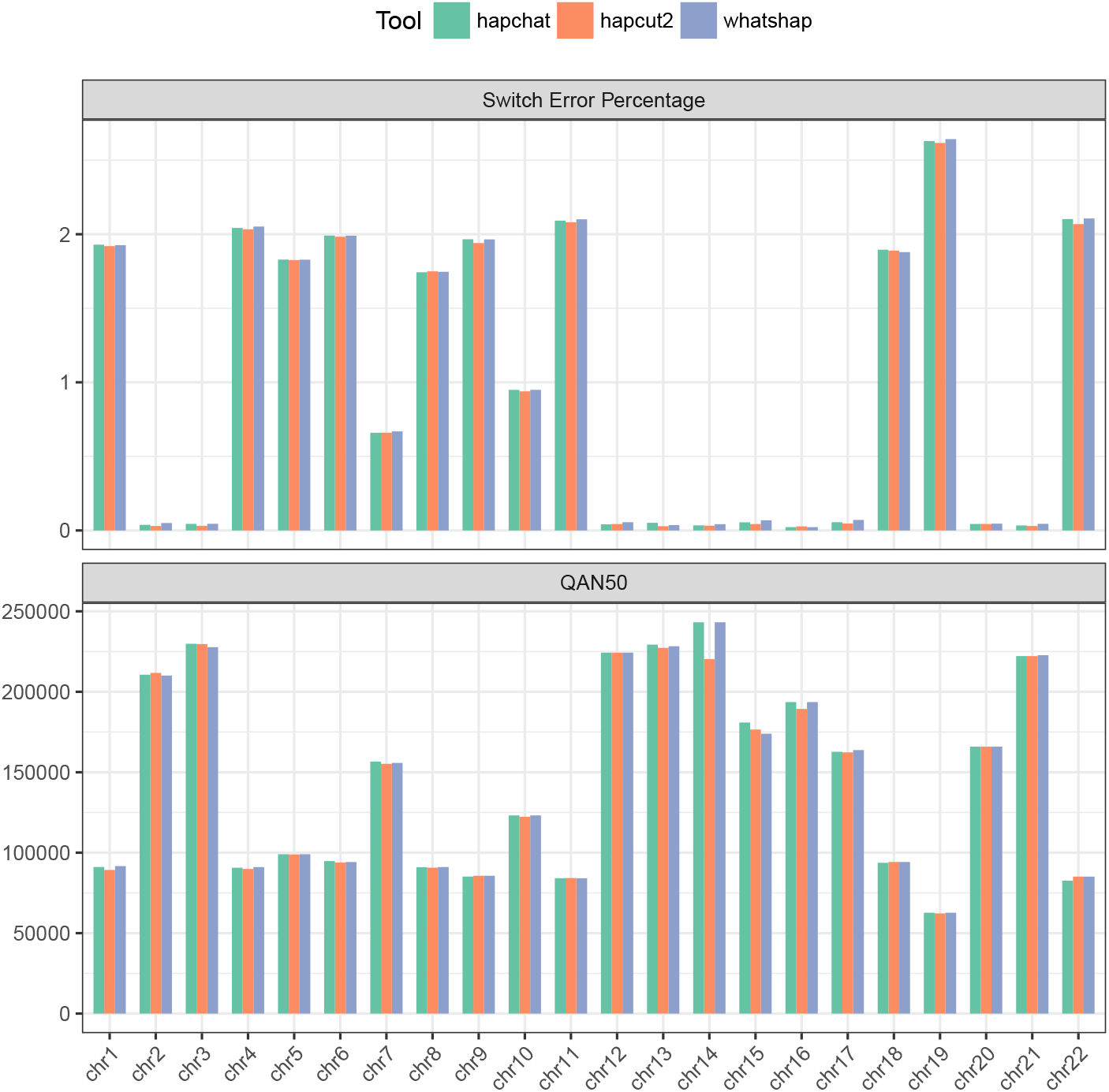
Quality measures on the real NA12878 dataset. We present the bar plots showing the measures of switch error percentage and QAN50 achieved by HapCHAT, WhatsHap, and HapCUT2 on the different chromosome datasets of NA12878.

**Table 1.**
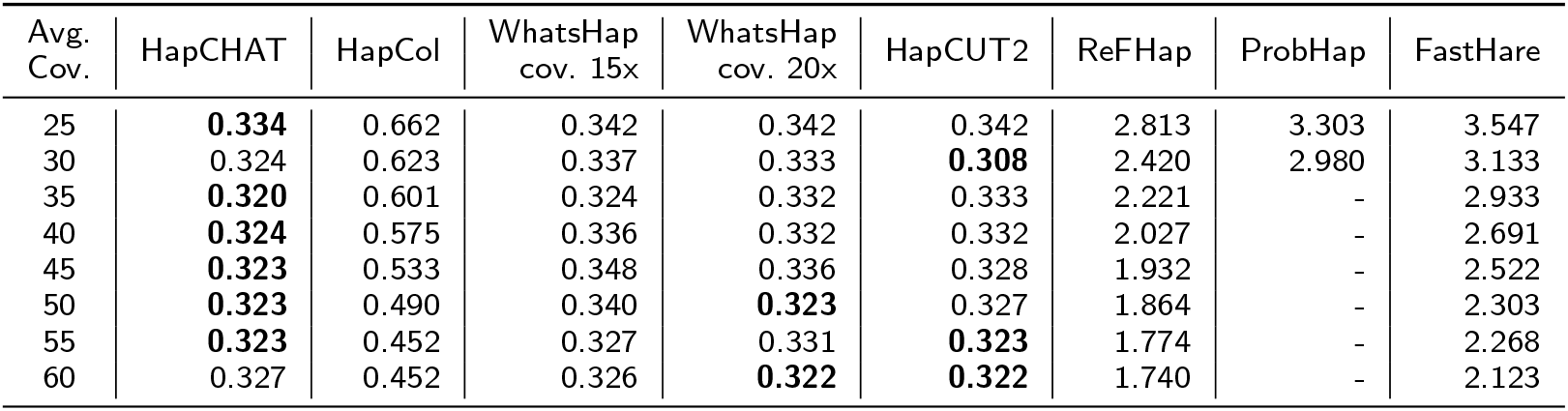
Switch error percentage on the real Ashkenazim dataset, Chromosome 1. For each dataset, its row identified by its average coverage (Avg. Cov.). We report the results obtained by running the tools with maximum coverage 30× for HapCHAT, 25× for HapCol, 15× and 20× for WhatsHap. No maximum coverage was set for HapCUT2, ReFHap, ProbHap, and FastHare. The best result (lowest value) for each dataset is boldfaced.

The accuracy of the predictions obtained from the experiments and measured in terms of switch error percentages is summarized in Tables 1, 6 and 11. We have also assessed the accuracy of the predictions by computing the Hamming distance percentages — Tables 2, 7 and 12. Each true haplotype is a mosaic of the predicted haplotypes. A switch error is the boundary (that is two consecutive SNV positions) between two portions of such a mosaic. The switch error percentage is the ratio between the number of switch errors and the number of phased SNVs minus one (expressed as a percentage). It is immediate to notice that HapCHAT, WhatsHap, and HapCUT2 compute the best predictions, all of them being very close. Figures 2 and 3 give bar chart representations of switch error rates for just these three methods on all real datasets. We point out that HapCHAT (resp., HapCUT2) computes the best switch error rates for almost all instances of the real and simulated Ashkenazim (resp., NA12878) datasets.

**Table 2.** Hamming distance on the real Ashkenazim dataset, Chromosome 1. For each dataset, its row identified by its average coverage (Avg. Cov.). We report the results obtained by running the tools with maximum coverage 30× for HapCHAT, 25× for HapCol, 15× and 20× for WhatsHap. No maximum coverage was set for HapCUT2, ReFHap, ProbHap, and FastHare. The best result (lowest value) for each dataset is boldfaced.

| Avg. Cov. | HapCHAT | HapCol | WhatsHap cov. 15x | WhatsHap cov. 20x | HapCUT2 | ReFHap | ProbHap | FastHare |
| --- | --- | --- | --- | --- | --- | --- | --- | --- |
| 25 | 0.54 | 2.41 | 0.64 | 0.84 | **0.44** | 3.96 | 3.42 | 5.53 |
| 30 | 0.35 | 2.18 | 0.64 | 0.60 | **0.24** | 3.46 | 3.41 | 5.38 |
| 35 | **0.36** | 2.02 | 0.37 | 0.42 | 0.37 | 3.99 | - | 5.62 |
| 40 | **0.37** | 1.66 | 0.45 | 0.44 | **0.37** | 3.10 | - | 5.08 |
| 45 | 0.38 | 1.80 | 0.43 | 0.42 | **0.37** | 3.02 | - | 4.49 |
| 50 | 0.41 | 1.47 | 0.41 | 0.38 | **0.35** | 2.84 | - | 4.32 |
| 55 | 0.40 | 0.87 | **0.36** | 0.41 | 0.37 | 3.28 | - | 4.67 |
| 60 | 0.39 | 1.25 | **0.34** | 0.36 | 0.35 | 3.60 | - | 5.06 |

Although the switch error is one of the most widely adopted measures used to evaluate the quality of the phased haplotypes, it does not take into account the *recall*, or the completeness of the haplotype – that is, the size of the phased haplotype blocks recovered. While N50 is the classical median size of an assembled haplotype block in terms of length in basepairs (bps) from the literature on assembly, [49] introduced the *adjusted* N50, that is AN50 score which normalizes each block in terms of the number of phased SNVs appearing on a block. In order to account for completeness *and* quality, [50] introduced the notion of *quality* AN50, that is the QAN50 score, where assembled haplotype blocks are fractured at each switch error, and then AN50 is taken on the resulting sub-blocks. This is an important measure because it is closest to the objective of haplotype assembly – to reassemble the longest (error-free) haplotype blocks possible. We hence computed QAN50 scores for all methods, as summarized in Tables 3, 8, and 13. It is immediate to notice that HapCHAT and WhatsHap have the best QAN50 scores, more precisely HapCHAT (resp., WhatsHap) computes the best QAN50 scores for almost all instances of the real and simulated Ashkenazim (resp., NA12878) datasets. HapCUT2 is a close second: despite its good switch error rate, it has consistently lower QAN50 scores.

**Table 3.**
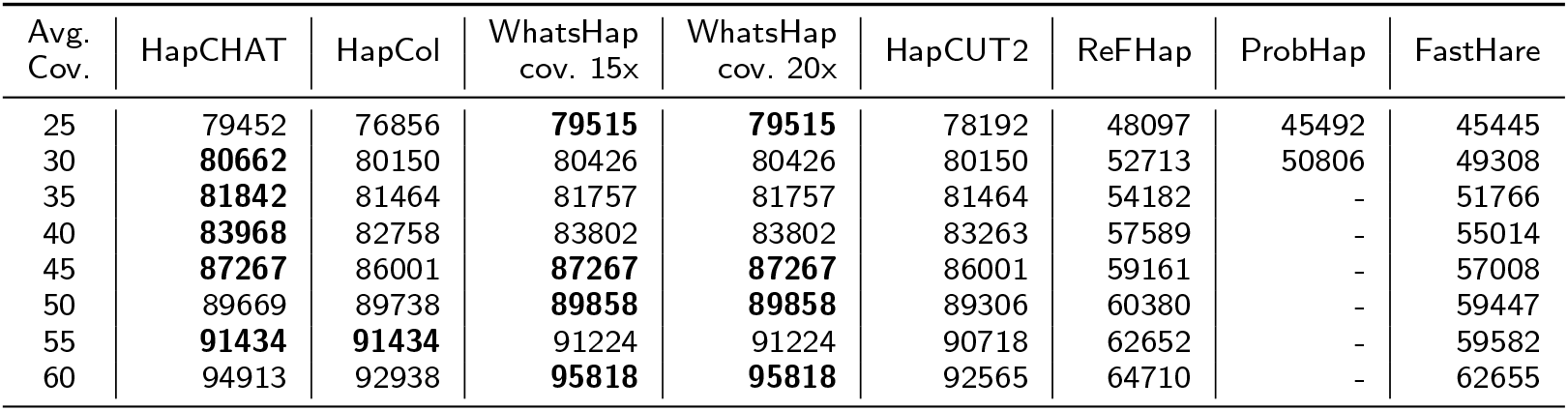
QAN50 on the real Ashkenazim dataset, Chromosome 1. For each dataset, its row identified by its average coverage (Avg. Cov.). We report the results obtained by running the tools with maximum coverage 30× for HapCHAT, 25× for HapCol, 15× and 20× for WhatsHap. No maximum coverage was set for HapCUT2, ReFHap, ProbHap, and FastHare. The best result (highest value) for each dataset is boldfaced.

| Avg. Cov. | HapCHAT | HapCol | WhatsHap cov. 15x | WhatsHap cov. 20x | HapCUT2 | ReFHap | ProbHap | FastHare |
| --- | --- | --- | --- | --- | --- | --- | --- | --- |
| 25 | 79452 | 76856 | **79515** | **79515** | 78192 | 48097 | 45492 | 45445 |
| 30 | **80662** | 80150 | 80426 | 80426 | 80150 | 52713 | 50806 | 49308 |
| 35 | **81842** | 81464 | 81757 | 81757 | 81464 | 54182 | - | 51766 |
| 40 | **83968** | 82758 | 83802 | 83802 | 83263 | 57589 | - | 55014 |
| 45 | **87267** | 86001 | **87267** | **87267** | 86001 | 59161 | - | 57008 |
| 50 | 89669 | 89738 | **89858** | **89858** | 89306 | 60380 | - | 59447 |
| 55 | **91434** | **91434** | 91224 | 91224 | 90718 | 62652 | - | 59582 |
| 60 | 94913 | 92938 | **95818** | **95818** | 92565 | 64710 | - | 62655 |

This could possibly be explained by [17]: “HapCUT2 implements likelihood-based strategies for pruning low-confidence variants to reduce mismatch errors and splitting blocks at poor linkages to reduce switch errors (see Methods). These postprocessing steps allow a user to improve accuracy of the haplotypes at the cost of reducing completeness and contiguity.” – indeed their switch error rate tends to be consistently the best for the NA12878 dataset at least, the tradeoff being that QAN50 score is consistently lower than the best method in all cases. Figures 2 and 3 give bar chart representations of QAN50 scores for HapCHAT, WhatsHap and HapCUT2 on all real datasets.

Since HapCHAT and WhatsHap can be influenced by a maximum coverage parameter, we did a deeper analysis of these two methods at different values of such parameter. The plots in Figure 1 represent the quality of the predictions computed by WhatsHap and HapCHAT as a function of the running time, for Chromosome 1 on the Ashkenazim dataset. Besides the switch error rate, we have also investigated the Hamming distance, that is the number of phase-calls that are different from the ground truth. Both plots confirm that HapCHAT computes predictions that are at least as good as those of WhatsHap (and clearly better in terms of Hamming distance) with a comparable runtime. We decided to include in the Tables the comparison of WhatsHap at both 20x and 15x max coverage, while 20x is the maximum coverage that we could test for WhatsHap – 15x is suggested by the authors as the default value for running WhatsHap and achieve the best trade off between accuracy and running time [48]. Observe in Figure 1 that with 20x max coverage WhatsHap obtains better predictions — close to those by HapCHAT — but with a much higher runtime.

It is possible to observe from Tables 4, 5, 9 and 10 that although both time and memory used by HapCHAT is growing with the (average) coverage, with higher coverage the rate at which the time increments is decreasing. Similarly, also the memory increment is almost linear with respect to the growth of the coverage of the datasets. On the other hand, while the changes of time and memory required by HapCol and WhatsHap to process higher coverages remain similar. Contrary to HapCHAT, because HapCol and WhatsHap are not adaptive (see intro for more details) that is they do not change their behaviour w.r.t. increasing average coverage, they must be run at a uniform *maximum* coverage of 25 and 15, respectively, and exhibit similar runtimes and memory usage for all datasets. HapCHAT, on the other hand, processes these datasets at the higher uniform maximum coverage of 30, and because it *adapts* to this increased average coverage, we see this linear trend in increased resource usage, as expected. Finally, we point out that HapCUT2, ReFHap, and FastHare require always the same memory, since it does not depend on the coverage, and the time grows linearly, while ProbHap exhibits a behavior reflecting the coverage increment, especially in terms of memory consumption.

**Table 4.** Time in seconds of the tools on real Ashkenazim datasets of Chromosome 1. For each dataset, its row identified by its average coverage (Avg. Cov.). We report the results obtained by running the tools with maximum coverage 30× for HapCHAT, 25× for HapCol, 15× and 20× for WhatsHap. No maximum coverage was set for HapCUT2, ReFHap, ProbHap, and FastHare.

| Avg. Cov. | HapCHAT | HapCol | WhatsHap cov. 15x | WhatsHap cov. 20x | HapCUT2 | ReFHap | ProbHap | FastHare |
| --- | --- | --- | --- | --- | --- | --- | --- | --- |
| 25 | 591 | 39456 | 1115 | 9278 | 1563 | 80 | 43573 | 3 |
| 30 | 1292 | 46564 | 1031 | 10753 | 1596 | 196 | 72696 | 4 |
| 35 | 2193 | 50071 | 1122 | 11959 | 1888 | 308 | - | 4 |
| 40 | 3095 | 50301 | 1247 | 12570 | 2160 | 499 | - | 5 |
| 45 | 3888 | 51570 | 1308 | 12735 | 2388 | 822 | - | 6 |
| 50 | 4579 | 53030 | 1395 | 12996 | 2731 | 1192 | - | 8 |
| 55 | 5103 | 54012 | 1534 | 13252 | 2983 | 1777 | - | 9 |
| 60 | 5550 | 53496 | 1605 | 13469 | 3216 | 2493 | - | 13 |

**Table 5.** Peak of RAM usage in Megabytes of the tools on real Ashkenazim datasets of Chromosome 1. For each dataset, its row identified by its average coverage (Avg. Cov.). We report the results obtained by running the tools with maximum coverage 30× for HapCHAT, 25× for HapCol, 15× and 20× for WhatsHap. No maximum coverage was set for HapCUT2, ReFHap, ProbHap, and FastHare.

| Avg. Cov. | HapCHAT | HapCol | WhatsHap cov. 15x | WhatsHap cov. 20x | HapCUT2 | ReFHap | ProbHap | FastHare |
| --- | --- | --- | --- | --- | --- | --- | --- | --- |
| 25 | 1370 | 2263 | 930 | 5510 | 3266 | 3005 | 4693 | 3005 |
| 30 | 1661 | 2562 | 931 | 6195 | 3270 | 3005 | 5355 | 3005 |
| 35 | 1966 | 2908 | 931 | 6513 | 3276 | 3005 | - | 3005 |
| 40 | 2291 | 3231 | 931 | 6483 | 3279 | 3005 | - | 3005 |
| 45 | 2636 | 3190 | 952 | 6937 | 3283 | 3005 | - | 3005 |
| 50 | 3158 | 3286 | 1007 | 7144 | 3287 | 3005 | - | 3005 |
| 55 | 3549 | 3479 | 1042 | 7229 | 3292 | 3005 | - | 3005 |
| 60 | 3968 | 5412 | 1073 | 7430 | 3296 | 3005 | - | 3005 |

An analysis of Tables 1 and 6 towards finding the effect of average coverage shows that there is a trend of improving predictions with higher average coverage, but this improvement is irregular. Since those irregularities are more common for HapCHAT than for the other tools, we have produced Table 17 which gives a more detailed breakdown of how the switch error is changing as a result of increasing coverage. More precisely, we have found that only in one case the erroneous sites at higher coverage is a subset of the erroneous sites at lower coverage. This shows a higher sensitivity of HapCHAT to changing (in this case sampling) instances. On the other hand, the quality measure given by the QAN50 reported in Tables 3 and 8 and also summarized in Figure 2 shows that there is a regular increase of the QAN50 for all the data sets consistent with the increase of the coverage.

**Table 6.** Switch error percentage on simulated datasets of Chromosome 1. For each dataset, its row identified by its average coverage (Avg. Cov.). We report the results obtained by running the tools with maximum coverage 30× for HapCHAT, 25× for HapCol, 15× and 20× for WhatsHap. No maximum coverage was set for HapCUT2, ReFHap, ProbHap, and FastHare. The best result (lowest value) for each dataset is boldfaced.

| Avg. Cov. | HapCHAT | HapCol | WhatsHap cov. 15x | WhatsHap cov. 20x | HapCUT2 | ReFHap | ProbHap | FastHare |
| --- | --- | --- | --- | --- | --- | --- | --- | --- |
| 25 | **0.035** | 0.218 | **0.035** | 0.039 | 0.037 | 1.081 | 1.487 | 2.112 |
| 30 | **0.028** | 0.181 | 0.035 | 0.031 | 0.037 | 0.725 | 1.166 | 1.430 |
| 35 | **0.028** | 0.161 | 0.033 | 0.037 | 0.037 | 0.537 | 0.879 | 1.086 |
| 40 | **0.026** | 0.148 | **0.026** | 0.030 | 0.037 | 0.425 | - | 0.901 |
| 45 | **0.022** | 0.139 | 0.024 | 0.024 | **0.022** | 0.404 | - | 0.781 |
| 50 | 0.020 | 0.134 | 0.020 | 0.024 | **0.018** | 0.324 | - | 0.586 |
| 55 | 0.022 | 0.126 | 0.024 | 0.022 | **0.018** | 0.273 | - | 0.565 |
| 60 | **0.020** | 0.108 | **0.020** | 0.024 | 0.022 | 0.248 | - | 0.470 |

**Table 7.** Hamming Distance percentage on simulated datasets of Chromosome 1. For each dataset, its row identified by its average coverage (Avg. Cov.). We report the results obtained by running the tools with maximum coverage 30× for HapCHAT, 25× for HapCol, 15× and 20× for WhatsHap. No maximum coverage was set for HapCUT2, ReFHap, ProbHap, and FastHare. The best result (lowest value) for each dataset is boldfaced.

| Avg. Cov. | HapCHAT | HapCol | WhatsHap cov. 15x | WhatsHap cov. 20x | HapCUT2 | ReFHap | ProbHap | FastHare |
| --- | --- | --- | --- | --- | --- | --- | --- | --- |
| 25 | 0.35 | 1.30 | 0.42 | 0.42 | **0.33** | 3.43 | 2.66 | 5.95 |
| 30 | **0.33** | 1.92 | 0.43 | 0.42 | 0.38 | 2.52 | 2.43 | 4.55 |
| 35 | **0.27** | 1.37 | 0.34 | 0.55 | 0.37 | 1.92 | 2.09 | 3.93 |
| 40 | 0.26 | 1.18 | **0.24** | 0.41 | 0.27 | 1.86 | - | 3.31 |
| 45 | 0.34 | 1.02 | 0.32 | 0.34 | **0.27** | 1.95 | - | 3.12 |
| 50 | **0.27** | 1.18 | 0.78 | 0.81 | 0.73 | 1.42 | - | 3.14 |
| 55 | 0.28 | 1.13 | 0.76 | 0.76 | **0.20** | 1.48 | - | 3.19 |
| 60 | **0.13** | 1.26 | 0.17 | 0.16 | 0.57 | 1.49 | - | 3.49 |

**Table 8.** QAN50 results of the tools on real simulated datasets of Chromosome 1. For each dataset, its row identified by its average coverage (Avg. Cov.). We report the results obtained by running the tools with maximum coverage 30× for HapCHAT, 25× for HapCol, 15× and 20× for WhatsHap. No maximum coverage was set for HapCUT2, ReFHap, ProbHap, and FastHare. The best result (highest value) for each dataset is boldfaced.

| Avg. Cov. | HapCHAT | HapCol | WhatsHap cov. 15x | WhatsHap cov. 20x | HapCUT2 | ReFHap | ProbHap | FastHare |
| --- | --- | --- | --- | --- | --- | --- | --- | --- |
| 25 | **87890** | 85002 | 87581 | 87581 | 51183 | 50325 | 49121 | 45846 |
| 30 | **93454** | 87599 | 92831 | 92565 | 57323 | 56745 | 54412 | 52138 |
| 35 | **96311** | 92483 | 96167 | 95611 | 61204 | 60612 | 59047 | 56881 |
| 40 | **97810** | 95818 | **97810** | 97270 | 64979 | 64535 | - | 60748 |
| 45 | **100826** | 98674 | **100826** | **100826** | 68274 | 66973 | - | 64003 |
| 50 | **103348** | 100826 | **103348** | **103348** | 73159 | 73256 | - | 69457 |
| 55 | 105243 | 103348 | **106341** | **106341** | 74273 | 74402 | - | 71058 |
| 60 | 107121 | 105243 | **107569** | **107569** | 76497 | 76497 | - | 73256 |

**Table 9.** Time in seconds of the tools on simulated datasets of Chromosome 1. For each dataset, its row identified by its average coverage (Avg. Cov.). We report the results obtained by running the tools with maximum coverage 30× for HapCHAT, 25× for HapCol, 15× and 20× for WhatsHap. No maximum coverage was set for HapCUT2, ReFHap, ProbHap, and FastHare.

| Avg. Cov. | HapCHAT | HapCol | WhatsHap cov. 15x | WhatsHap cov. 20x | HapCUT2 | ReFHap | ProbHap | FastHare |
| --- | --- | --- | --- | --- | --- | --- | --- | --- |
| 25 | 572 | 38863 | 1027 | 9686 | 205 | 44 | 33988 | 3 |
| 30 | 1317 | 47367 | 883 | 11095 | 238 | 91 | 56165 | 3 |
| 35 | 2167 | 18813 | 954 | 11650 | 286 | 167 | 80061 | 4 |
| 40 | 3052 | 20007 | 1048 | 12760 | 323 | 269 | - | 5 |
| 45 | 3754 | 56403 | 1161 | 12678 | 367 | 423 | - | 6 |
| 50 | 4399 | 57135 | 1170 | 12860 | 412 | 672 | - | 6 |
| 55 | 4882 | 56745 | 1287 | 13174 | 467 | 1019 | - | 7 |
| 60 | 5277 | 21070 | 1336 | 13407 | 496 | 1536 | - | 9 |

**Table 10.**
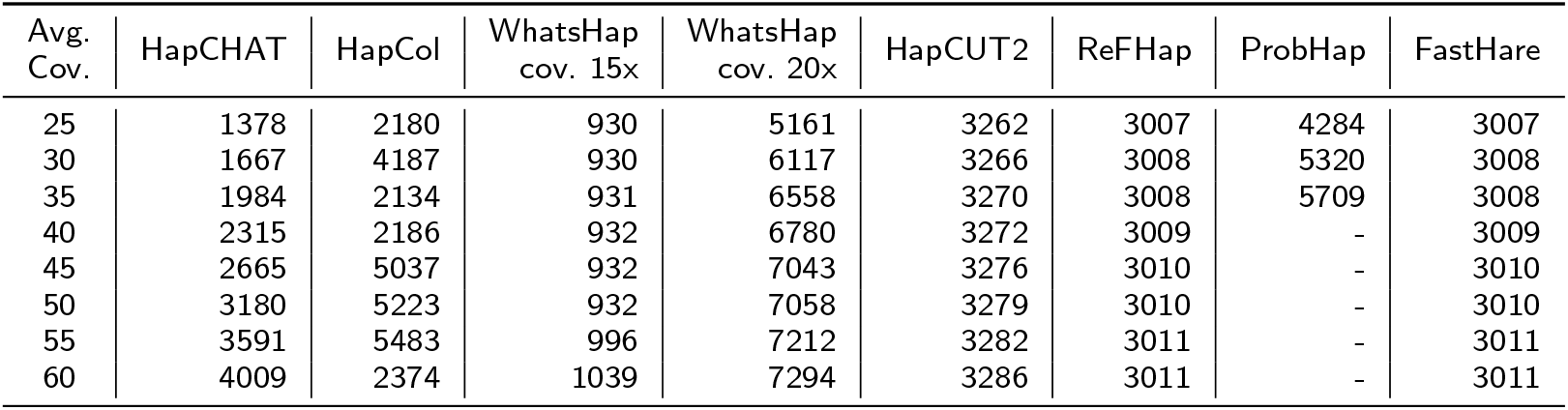
Peak of RAM usage in Megabytes of the tools on simulated datasets of Chromosome 1. For each dataset, its row identified by its average coverage (Avg. Cov.). We report the results obtained by running the tools with maximum coverage 30× for HapCHAT, 25× for HapCol, 15× and 20× for WhatsHap. No maximum coverage was set for HapCUT2, ReFHap, ProbHap, and FastHare.

**Table 11.** Switch error percentage on datasets of NA12878. Each row corresponds to a chromosome. The dataset consists of all reads aligned to the chromosome. We report the results obtained by running the tools with maximum coverage 30× for HapCHAT, 25× for HapCol, 15× and 20× for WhatsHap. No maximum coverage was set for HapCUT2, ReFHap, ProbHap, and FastHare. The best result (lowest value) for each dataset is boldfaced.

| Chrom. | HapCHAT | HapCol | WhatsHap<br>cov. 15x | WhatsHap<br>cov. 20x | HapCUT2 | ReFHap | ProbHap | FastHare |
| --- | --- | --- | --- | --- | --- | --- | --- | --- |
| 1 | 1.929 | - | 1.926 | 1.924 | **1.920** | - | - | 2.191 |
| 2 | 0.038 | - | 0.050 | 0.035 | **0.030** | - | - | 0.374 |
| 3 | 0.044 | - | 0.045 | 0.039 | **0.031** | - | - | 0.381 |
| 4 | 2.042 | - | 2.052 | 2.048 | **2.033** | - | - | 2.237 |
| 5 | 1.829 | - | 1.828 | **1.824** | 1.825 | - | - | 1.998 |
| 6 | 1.991 | - | 1.990 | 1.991 | **1.983** | - | - | 2.205 |
| 7 | **0.659** | - | 0.669 | 0.666 | 0.660 | - | - | 0.924 |
| 8 | **1.743** | - | 1.746 | 1.748 | 1.749 | - | - | 1.992 |
| 9 | 1.966 | - | 1.965 | 1.966 | **1.940** | 2.140 | - | 2.187 |
| 10 | 0.949 | - | 0.949 | 0.948 | **0.939** | 1.171 | - | 1.232 |
| 11 | 2.092 | - | 2.101 | 2.101 | **2.081** | 2.282 | - | 2.325 |
| 12 | **0.041** | - | 0.055 | 0.048 | 0.043 | 0.319 | - | 0.405 |
| 13 | 0.051 | - | 0.036 | 0.049 | **0.029** | 0.285 | - | 0.349 |
| 14 | 0.034 | - | 0.042 | 0.039 | **0.032** | 0.347 | - | 0.421 |
| 15 | 0.055 | 0.331 | 0.069 | 0.065 | **0.043** | 0.358 | - | 0.427 |
| 16 | **0.022** | 0.289 | **0.022** | 0.029 | 0.027 | 0.322 | - | 0.420 |
| 17 | 0.055 | 0.277 | 0.071 | 0.067 | **0.047** | 0.337 | - | 0.426 |
| 18 | 1.895 | - | 1.879 | **1.876** | 1.889 | 2.072 | - | 2.122 |
| 19 | 2.629 | - | 2.642 | 2.644 | **2.616** | 2.807 | - | 2.914 |
| 20 | **0.043** | 0.277 | 0.046 | **0.043** | **0.043** | 0.412 | - | 0.451 |
| 21 | 0.033 | - | 0.044 | 0.041 | **0.030** | 0.364 | - | 0.408 |
| 22 | 2.102 | 2.323 | 2.106 | 2.114 | **2.068** | 2.378 | - | 2.452 |

**Table 12.** Hamming Distance percentage on datasets of NA12878. Each row corresponds to a chromosome. The dataset consists of all reads aligned to the chromosome. We report the results obtained by running the tools with maximum coverage 30× for HapCHAT, 25× for HapCol, 15× and 20× for WhatsHap. No maximum coverage was set for HapCUT2, ReFHap, ProbHap, and FastHare. The best result (lowest value) for each dataset is boldfaced.

| Chrom. | HapCHAT | HapCol | WhatsHap<br>cov. 15x | WhatsHap<br>cov. 20x | HapCUT2 | ReFHap | ProbHap | FastHare |
| --- | --- | --- | --- | --- | --- | --- | --- | --- |
| 1 | 2.12 | - | **1.92** | 2.10 | 2.11 | - | - | 6.16 |
| 2 | 0.51 | - | 0.49 | **0.29** | 0.77 | - | - | 4.91 |
| 3 | **0.32** | - | 0.42 | 0.42 | 0.48 | - | - | 4.74 |
| 4 | 2.47 | - | 2.15 | 2.18 | **2.00** | - | - | 6.44 |
| 5 | 2.33 | - | 2.53 | 2.22 | **1.98** | - | - | 6.56 |
| 6 | 3.39 | - | 3.02 | 3.20 | **2.82** | - | - | 7.15 |
| 7 | 1.16 | - | **1.10** | **1.10** | 1.36 | - | - | 5.05 |
| 8 | 2.44 | - | 2.46 | 2.54 | **2.02** | - | - | 6.14 |
| 9 | 2.45 | - | 2.31 | 2.49 | **2.11** | 5.68 | - | 6.23 |
| 10 | 1.19 | - | 1.16 | 1.18 | **0.93** | 3.89 | - | 5.29 |
| 11 | 2.08 | - | 2.06 | 2.06 | **1.99** | 4.25 | - | 5.08 |
| 12 | 0.43 | - | 0.48 | **0.38** | 0.51 | 2.92 | - | 5.54 |
| 13 | 0.41 | - | 0.63 | 0.57 | **0.35** | 4.01 | - | 4.84 |
| 14 | 0.21 | - | 0.48 | 0.58 | **0.17** | 3.01 | - | 3.24 |
| 15 | **0.23** | 3.39 | 0.24 | 0.34 | 0.34 | 4.18 | - | 5.49 |
| 16 | **0.24** | 2.09 | 0.45 | 0.88 | 0.28 | 1.65 | - | 2.87 |
| 17 | 0.50 | 2.84 | 0.38 | 0.79 | **0.20** | 2.89 | - | 4.61 |
| 18 | 1.80 | - | 1.67 | **1.65** | 1.68 | 4.77 | - | 8.10 |
| 19 | 3.19 | - | 3.14 | 3.40 | **2.99** | 4.37 | - | 7.32 |
| 20 | 1.37 | 3.47 | 0.16 | **0.10** | **0.16** | 2.99 | - | 4.07 |
| 21 | **0.10** | - | **0.10** | **0.10** | 1.95 | 5.37 | - | 4.22 |
| 22 | **1.82** | 4.92 | 1.84 | 1.83 | **1.82** | 4.83 | - | 6.52 |

**Table 13.** QAN50 results of the tools on datasets of NA12878. Each row corresponds to a chromosome. The dataset consists of all reads aligned to the chromosome. We report the results obtained by running the tools with maximum coverage 30× for HapCHAT, 25× for HapCol, 15× and 20× for WhatsHap. No maximum coverage was set for HapCUT2, ReFHap, ProbHap, and FastHare. The best result (lowest value) for each dataset is boldfaced.

| Chrom. | HapCHAT | HapCol | WhatsHap<br>cov. 15x | WhatsHap<br>cov. 20x | HapCUT2 | ReFHap | ProbHap | FastHare |
| --- | --- | --- | --- | --- | --- | --- | --- | --- |
| 1 | 91098 | - | 91668 | **91677** | 89249 | - | - | 84863 |
| 2 | 210603 | - | 210098 | **211732** | **211732** | - | - | 177388 |
| 3 | **229835** | - | 227732 | **229835** | 229655 | - | - | 170494 |
| 4 | 90639 | - | **91034** | 90639 | 89868 | - | - | 84861 |
| 5 | 99011 | - | 99012 | **99567** | 98900 | - | - | 91745 |
| 6 | **94780** | - | 94200 | **94780** | 93894 | - | - | 85483 |
| 7 | **156573** | - | 155773 | 155773 | 155209 | - | - | 135095 |
| 8 | 90928 | - | **91069** | 90836 | 90661 | - | - | 84076 |
| 9 | 85172 | - | **85655** | 85469 | **85655** | 82917 | - | 80957 |
| 10 | 123171 | - | 123171 | **123224** | 122317 | 114172 | - | 112861 |
| 11 | 84153 | - | 84108 | 84108 | **84237** | 81526 | - | 79057 |
| 12 | 224308 | - | 224308 | **228356** | 224308 | 190161 | - | 174540 |
| 13 | **229318** | - | 228310 | 228310 | 227286 | 178173 | - | 175124 |
| 14 | **243192** | - | **243192** | 227040 | 220294 | 186476 | - | 181826 |
| 15 | **180874** | 153527 | 173950 | 176529 | 176529 | 147339 | - | 138185 |
| 16 | **193611** | 160049 | **193611** | 190884 | 189342 | 158848 | - | 152960 |
| 17 | 162690 | 151262 | **163789** | **163789** | 162328 | 140216 | - | 133887 |
| 18 | 93705 | - | **94210** | 93705 | **94210** | 87076 | - | 83383 |
| 19 | **62662** | - | **62662** | 62568 | 62233 | 59716 | - | 58694 |
| 20 | 165921 | 163062 | 165921 | **176807** | 165921 | 140498 | - | 140034 |
| 21 | 222171 | - | **222769** | 221786 | 222171 | 149165 | - | 146675 |
| 22 | 82618 | 73223 | **85112** | 82618 | **85112** | 72117 | - | 70718 |

**Table 14.** Time in seconds on datasets of NA12878. Each row corresponds to a chromosome. The dataset consists of all reads aligned to the chromosome. We report the results obtained by running the tools with maximum coverage 30× for HapCHAT, 25× for HapCol, 15× and 20× for WhatsHap. No maximum coverage was set for HapCUT2, ReFHap, ProbHap, and FastHare.

| Chrom. | HapCHAT | HapCol | WhatsHap<br>cov. 15x | WhatsHap<br>cov. 20x | HapCUT2 | ReFHap | ProbHap | FastHare |
| --- | --- | --- | --- | --- | --- | --- | --- | --- |
| 1 | 20183 | - | 3300 | 41626 | 6301 | - | - | 980 |
| 2 | 21913 | - | 3686 | 46937 | 6758 | - | - | 1075 |
| 3 | 19325 | - | 2994 | 38040 | 5536 | - | - | 776 |
| 4 | 21744 | - | 3031 | 40083 | 5998 | - | - | 862 |
| 5 | 18416 | - | 2943 | 36674 | 5169 | - | - | 790 |
| 6 | 17792 | - | 2658 | 35189 | 5640 | - | - | 759 |
| 7 | 14321 | - | 2409 | 32550 | 4429 | - | - | 744 |
| 8 | 15930 | - | 2421 | 29902 | 4578 | - | - | 669 |
| 9 | 11307 | - | 1886 | 23586 | 3369 | 86913 | - | 635 |
| 10 | 13943 | - | 2244 | 27638 | 3914 | 86941 | - | 670 |
| 11 | 13291 | - | 1983 | 25419 | 3916 | 86833 | - | 567 |
| 12 | 12684 | - | 1916 | 25865 | 4054 | 86814 | - | 554 |
| 13 | 11100 | - | 1474 | 20288 | 2952 | 86686 | - | 406 |
| 14 | 9017 | - | 1265 | 17658 | 2644 | 86684 | - | 384 |
| 15 | 6934 | 63221 | 1114 | 14218 | 2102 | 86700 | - | 368 |
| 16 | 7426 | 69771 | 1265 | 16323 | 2589 | 86783 | - | 461 |
| 17 | 6460 | 54037 | 956 | 12312 | 1832 | 86669 | - | 312 |
| 18 | 8440 | - | 1152 | 15794 | 2497 | 86671 | - | 353 |
| 19 | 3625 | - | 826 | 10368 | 1617 | 86668 | - | 296 |
| 20 | 5878 | 55032 | 827 | 11815 | 1594 | 86600 | - | 243 |
| 21 | 3561 | - | 560 | 7508 | 1308 | 86585 | - | 226 |
| 22 | 2835 | 31617 | 505 | 7059 | 1002 | 86568 | - | 195 |

**Table 15.** Peak of RAM usage in Megabytes of the tools on datasets of NA12878. Each row corresponds to a chromosome. The dataset consists of all reads aligned to the chromosome. We report the results obtained by running the tools with maximum coverage 30× for HapCHAT, 25× for HapCol, 15× and 20× for WhatsHap. No maximum coverage was set for HapCUT2, ReFHap, ProbHap, and FastHare.

| Chrom. | HapCHAT | HapCol | WhatsHap<br>cov. 15x | WhatsHap<br>cov. 20x | HapCUT2 | ReFHap | ProbHap | FastHare |
| --- | --- | --- | --- | --- | --- | --- | --- | --- |
| 1 | 6361 | - | 2983 | 13259 | 3351 | - | - | 3050 |
| 2 | 7082 | - | 3173 | 13938 | 3362 | - | - | 3056 |
| 3 | 6180 | - | 2669 | 12672 | 3329 | - | - | 3041 |
| 4 | 7531 | - | 2685 | 12959 | 3334 | - | - | 3046 |
| 5 | 5882 | - | 2551 | 12364 | 3320 | - | - | 3033 |
| 6 | 5649 | - | 2325 | 12120 | 3312 | - | - | 3031 |
| 7 | 4597 | - | 2080 | 11167 | 3309 | - | - | 3022 |
| 8 | 5075 | - | 2091 | 11164 | 3302 | - | - | 3023 |
| 9 | 3583 | - | 1639 | 9345 | 3285 | 17915 | - | 3009 |
| 10 | 4059 | - | 1819 | 10164 | 3303 | 9766 | - | 3017 |
| 11 | 3965 | - | 1814 | 10135 | 3290 | 9632 | - | 3018 |
| 12 | 4011 | - | 1787 | 10229 | 3288 | 13984 | - | 3016 |
| 13 | 7950 | - | 1449 | 8982 | 3267 | 8371 | - | 3006 |
| 14 | 2857 | - | 1281 | 8198 | 3261 | 10024 | - | 2998 |
| 15 | 2232 | 8437 | 1077 | 7370 | 3257 | 8302 | - | 2993 |
| 16 | 3116 | 19698 | 1128 | 7703 | 3263 | 10328 | - | 2995 |
| 17 | 7844 | 7845 | 962 | 6737 | 3253 | 5941 | - | 2990 |
| 18 | 3542 | - | 1152 | 7810 | 3254 | 15868 | - | 2995 |
| 19 | 1721 | - | 793 | 6055 | 3244 | 8808 | - | 2983 |
| 20 | 8496 | 9966 | 865 | 6612 | 3242 | 7973 | - | 2985 |
| 21 | 1329 | - | 611 | 5211 | 3229 | 7852 | - | 2977 |
| 22 | 3324 | 7782 | 542 | 4912 | 3225 | 7904 | - | 2975 |

Table 16 reports for each of the 16 Ashkenazim datasets, the SNV sites when the adaptive procedure of subsection Adaptive *k*-cMEC was activated. Interestingly, it is only in the Simulated dataset that the number of corrections needed to be increased from 5 to 8 – the rest needing an increase only to 7 (from 4) – indicating that it contains more unanticipated errors than the real datasets. Indeed this demonstrates that this adaptive procedure is an improvement over HapCol, recalling that each time this procedure is invoked, HapCol fails by definition. An added benefit of this procedure is that it can serve as an indicator of the quality of the read set to be phased. More specifically, it can serve as an indicator of the quality of the variant calling itself — indeed it is a third type of accuracy prediction, on top of switch error and Hamming distance — one that can be used to integrate the predictions of several tools to obtain higher quality variant calls [41, 42]. We plan to investigate further this advantage in future developments of HapCHAT.

**Table 16.** List of SNV positions when the adaptive procedure of subsection Adaptive *k*-cMEC was activated for real Ashkenazim and simulated datasets of Chromosome 1. For each dataset, its row is identified by its average coverage (Avg. Cov.). The positions in column ‘4 to 7’ are those for which the number of corrections was increased from 4 to 7, and similarly for the column ‘5 to 8’.

| Chr.1 |  | 4 to 7 |  | 5 to 8 |
| --- | --- | --- | --- | --- |
| Data | Avg. Cov. |  |  |  |
| Ashkenazim | Cov. 25 |  |  |  |
|  | Cov. 30 |  |  |  |
|  | Cov. 35 | 35556 |  |  |
|  | Cov. 40 |  |  |  |
|  | Cov. 45 | 35581 |  |  |
|  | Cov. 50 | 35593, 42897 |  |  |
|  | Cov. 55 | 3528 |  |  |
|  | Cov. 60 | 46338, 46339 |  |  |
| Simulated | Cov. 25 | 35569, 38788 |  | 26778 |
|  | Cov. 30 | 35594, 38815, 38817 |  | 26800 |
|  | Cov. 35 | 38827 |  | 26811 |
|  | Cov. 40 | 38837 |  | 38834, 38835, 38836 |
|  | Cov. 45 | 38844 |  | 38842 |
|  | Cov. 50 |  |  | 38849 |
|  | Cov. 55 |  |  |  |
|  | Cov. 60 |  |  |  |

**Table 17.** Comparison of the switch error positions on the Ashkenazim datasets of Chromosome 1 obtained with HapCHAT. For each pair of datasets having different coverages, we report the number of positions in which a switch error occurred as follows: those in common between the two datasets, those only found in the dataset of the row, and those only found the dataset of the column, respectively.

| Ashkenazim | Cov. 25 | Cov. 30 | Cov. 35 | Cov. 40 | Cov. 45 | Cov. 50 | Cov. 55 |
| --- | --- | --- | --- | --- | --- | --- | --- |
| Cov. 30 | 75/2/4 |  |  |  |  |  |  |
| Cov. 35 | 72/4/7 | 74/2/3 |  |  |  |  |  |
| Cov. 40 | 71/6/8 | 74/4/4 | 75/2/1 |  |  |  |  |
| Cov. 45 | 71/6/8 | 73/4/4 | 75/2/1 | 77/0/0 |  |  |  |
| Cov. 50 | 70/7/9 | 72/5/5 | 73/4/3 | 75/2/2 | 75/2/2 |  |  |
| Cov. 55 | 71/6/8 | 73/4/4 | 73/4/3 | 75/2/2 | 75/2/2 | 75/2/2 |  |
| Cov. 60 | 71/7/8 | 73/5/4 | 73/5/3 | 75/3/2 | 75/3/2 | 75/3/2 | 76/2/1 |

## Conclusions

We have presented HapCHAT, a tool that is able to phase high coverage PacBio reads. We have compared HapCHAT to WhatsHap, HapCol, HapCUT2, ReFHap, ProbHap and FastHare on on real and simulated whole-chromosome datasets, with average coverage up to 60×. The real datasets have been taken from the GIAB project. Our experimental comparison shows that HapCHAT has accuracy and recall that are comparable with those of WhatsHap and HapCUT2, and better than all other tools. At the same time, HapCHAT requires an amount of computational resources that is on the same order of magnitude as WhatsHap and HapCUT2. In particular, our QAN50 scores are almost consistently better than all other tools, showing that we reconstruct the longest, least fragmented haplotype blocks – the ultimate aim of haplotype assembly. Trying our dynamic programming approach with even *longer* reads, such as those bolstered with Hi-C information [51] would hence be an interesting future endeavour, to see how far we can push this method for assembling haplotypes.

Introducing the capability of adapting the number of errors permitted in each column allows HapCHAT to achieve a better fit than HapCol of the number of corrections needed at each variant site. Still, the current approach allows such adaptation only for the current column. Coupling this step with backtracking could result in fewer overall corrections.

Another direction of research is to fully consider the parent-sibling relations in trios, as done in [43] here. This is especially relevant, since most of the GIAB data is on trios.

Finally, we are working on the integration of HapCHAT with the WhatsHap tool to provide a more powerful haplotype phasing method able to combine the strengths of the two approaches.

## Declarations

### Ethics approval and consent to participate

Not applicable

### Consent for publication

Not applicable

### Availability of data and materials

The PacBio long reads data that we used are publicly available at ftp://ftp-trace.ncbi.nlm.nih.gov/giab/ftp/data/AshkenazimTrio/HG002_NA24385_son/PacBio_MtSinai_NIST/MtSinai_blasr_bam_GRCh37/hg002_gr37_1.bam and ftp://ftp-trace.ncbi.nlm.nih.gov/giab/ftp/data/NA12878/NA12878_PacBio_MtSinai/sorted_final_merged.bam The simulated datasets that we have used can be downloaded at: https://drive.google.com/drive/folders/0BxqLPsY2hmAXMlowZF9JQllZNEU. The BAM files are in `archiveHapCHAT-experiments_bam--simulated.tar.gz`, while the corresponding VCFs can be found at `HapCHAT-experiments.tar.gz`.

Instructions on how to use and replicate the experiments can be found at http://hapchat.algolab.eu

## Acknowledgments

We thank Tobias Marschall and Marcel Martin for inspiring discussions and for comments on earlier versions of this manuscript. We also thank the anonymous reviewers for pointing out during the revision process the new realignment feature of HapCut2 that allowed us to extend to experimental analysis and to use the QAN50 measure that helped the analysis and comparison of the tools.

## References

1. Browning, S.R., Browning, B.L.: Haplotype phasing: existing methods and new developments. Nature Reviews Genetics 12(10), 703–714 (2011)

2. Tewhey, R., Bansal, V., Torkamani, A., Topol, E.J., Schork, N.J.: The importance of phase information for human genomics. Nature Reviews Genetics (3), 215–223 (2011). doi:10.1038/nrg2950

3. Glusman, G., Cox, H.C., Roach, J.C.: Whole-genome haplotyping approaches and genomic medicine. Genome Medicine 6(9), 73 (2014). doi:10.1186/s13073-014-0073-7

4. Roach, J.C., Glusman, G., Smit, A.F.A., Huff, C.D., Hubley, R., Shannon, P.T., Rowen, L., Pant, K.P., Goodman, N., Bamshad, M., Shendure, J., Drmanac, R., Jorde, L.B., Hood, L., Galas, D.J.: Analysis of genetic inheritance in a family quartet by whole-genome sequencing. Science 328(5978), 636–639 (2010). doi:10.1126/science.1186802

5. Kuleshov, V., Xie, D., Chen, R., Pushkarev, D., Ma, Z., Blauwkamp, T., Kertesz, M., Snyder, M.: Whole-genome haplotyping using long reads and statistical methods. Nature Biotechnology 32(3), 261–266 (2014)

6. Porubský, D., Sanders, A.D., Wietmarschen, N.v., Falconer, E., Hills, M., Spierings, D.C.J., Bevova, M.R., Guryev, V., Lansdorp, P.M.: Direct chromosome-length haplotyping by single-cell sequencing. Genome Res. (2016)

7. Porubsky, D., Garg, S., Sanders, A.D., Korbel, J.O., Guryev, V., Lansdorp, P.M., Marschall, T.: Dense and accurate whole-chromosome haplotyping of individual genomes. Nat. Commun. 8(1), 1293 (2017)

8. Loh, P.-R., Danecek, P., Palamara, P.F., Fuchsberger, C., Reshef, Y.A., Finucane, H.K., Schoenherr, S., Forer, L., McCarthy, S., Abecasis, G.R., Durbin, R., Price, A.L.: Reference-based phasing using the haplotype reference consortium panel. Nature Genetics 48(11), 1443–1448 (2016). doi:10.1038/ng.3679

9. O’Connell, J., Sharp, K., Shrine, N., Wain, L., Hall, I., Tobin, M., Zagury, J.-F., Delaneau, O., Marchini, J.: Haplotype estimation for biobank-scale data sets. Nature Genetics 48(7), 817–820 (2016). doi:10.1038/ng.3583

10. Li, N., Stephens, M.: Modeling linkage disequilibrium and identifying recombination hotspots using single-nucleotide polymorphism data. Genetics 165(4), 2213–2233 (2003)

11. Lippert, R., Schwartz, R., Lancia, G., Istrail, S.: Algorithmic strategies for the single nucleotide polymorphism haplotype assembly problem. Briefings in Bioinformatics 3(1), 23–31 (2002)

12. Cilibrasi, R., Van Iersel, L., Kelk, S., Tromp, J.: The complexity of the single individual SNP haplotyping problem. Algorithmica 49(1), 13–36 (2007)

13. Bonizzoni, P., Dondi, R., Klau, G.W., Pirola, Y., Pisanti, N., Zaccaria, S.: On the fixed parameter tractability and approximability of the minimum error correction problem. In: 26th Annual Symposium on Combinatorial Pattern Matching (CPM). LNCS, vol. 9133, pp. 100–113 (2015)

14. Bonizzoni, P., Dondi, R., Klau, G.W., Pirola, Y., Pisanti, N., Zaccaria, S.: On the minimum error correction problem for haplotype assembly in diploid and polyploid genomes. Journal of Computational Biology 23(9), 718–736 (2016)

15. Panconesi, A., Sozio, M.: Fast hare: A fast heuristic for single individual SNP haplotype reconstruction. In: Algorithms in Bioinformatics, 4th International Workshop, WABI 2004, Bergen, Norway, September 17-21, 2004, Proceedings, pp. 266–277 (2004)

16. Bansal, V., Bafna, V.: HapCUT: an efficient and accurate algorithm for the haplotype assembly problem. Bioinformatics 24(16), 153–159 (2008)

17. Edge, P., Bafna, V., Bansal, V.: HapCUT2: robust and accurate haplotype assembly for diverse sequencing technologies. Genome Research 213462(116) (2016). doi:10.1101/gr.213462.116

18. Mazrouee, S., Wang, W.: FastHap: fast and accurate single individual haplotype reconstruction using fuzzy conflict graphs. Bioinformatics 30(17), 371–378 (2014)

19. Duitama, J., Huebsch, T., McEwen, G., Suk, E.-K., Hoehe, M.R.: ReFHap: a reliable and fast algorithm for single individual haplotyping. In: BCB, pp. 160–169 ACM, ??? (2010)

20. Fouilhoux, P., Mahjoub, A.R.: Solving VLSI design and DNA sequencing problems using bipartization of graphs. Computational Optimization and Applications 51(2), 749–781 (2012)

21. Chen, Z.-Z., Deng, F., Wang, L.: Exact algorithms for haplotype assembly from whole-genome sequence data. Bioinformatics 29(16), 1938–45 (2013)

22. He, D., Choi, A., Pipatsrisawat, K., Darwiche, A., Eskin, E.: Optimal algorithms for haplotype assembly from whole-genome sequence data. Bioinformatics 26(12), 183–190 (2010)

23. Chaisson, M.J.P., Sanders, A.D., Zhao, X., Malhotra, A., Porubsky, D., Rausch, T., Gardner, E.J., Rodriguez, O., Guo, L., Collins, R.L., Fan, X., Wen, J., Handsaker, R.E., Fairley, S., Kronenberg, Z.N., Kong, X., Hormozdiari, F., Lee, D., Wenger, A.M., Hastie, A., Antaki, D., Audano, P., Brand, H., Cantsilieris, S., Cao, H., Cerveira, E., Chen, C., Chen, X., Chin, C.-S., Chong, Z., Chuang, N.T., Church, D.M., Clarke, L., Farrell, A., Flores, J., Galeev, T., David, G., Gujral, M., Guryev, V., Haynes-Heaton, W., Korlach, J., Kumar, S., Kwon, J.Y., Lee, J.E., Lee, J., Lee, W.-P., Lee, S.P., Marks, P., Valud-Martinez, K., Meiers, S., Munson, K.M., Navarro, F., Nelson, B.J., Nodzak, C., Noor, A., Kyriazopoulou-Panagiotopoulou, S., Pang, A., Qiu, Y., Rosanio, G., Ryan, M., Stutz, A., Spierings, D.C.J., Ward, A., Welsch, A.E., Xiao, M., Xu, W., Zhang, C., Zhu, Q., Zheng-Bradley, X., Jun, G., Ding, L., Koh, C.L., Ren, B., Flicek, P., Chen, K., Gerstein, M.B., Kwok, P.-Y., Lansdorp, P.M., Marth, G., Sebat, J., Shi, X., Bashir, A., Ye, K., Devine, S.E., Talkowski, M., Mills, R.E., Marschall, T., Korbel, J., Eichler, E.E., Lee, C.: Multi-platform discovery of haplotype-resolved structural variation in human genomes. bioRxiv (2017). doi:10.1101/193144

24. Carneiro, M.O., Russ, C., Ross, M.G., Gabriel, S.B., Nusbaum, C., DePristo, M.A.: Pacific Biosciences sequencing technology for genotyping and variation discovery in human data. BMC Genomics 13(1), 375 (2012)

25. Roberts, R.J., Carneiro, M.O., Schatz, M.C.: The advantages of SMRT sequencing. Genome Biology 14(6), 405 (2013)

26. Sedlazeck, F.J., Lee, H., Darby, C.A., Schatz, M.C.: Piercing the dark matter: bioinformatics of long-range sequencing and mapping. Nat. Rev. Genet. (2018)

27. Kuleshov, V., et al. : Whole-genome haplotyping using long reads and statistical methods. Nature Biotechnology 32(3), 261–266 (2014)

28. Ip, C.L.C., Loose, M., Tyson, J.R., de Cesare, M., Brown, B.L., Jain, M., Leggett, R.M., Eccles, D.A., Zalunin, V., Urban, J.M., Piazza, P., Bowden, R.J., Paten, B., Mwaigwisya, S., Batty, E.M., Simpson, J.T., Snutch, T.P., Birney, E., Buck, D., Goodwin, S., Jansen, H.J., O’Grady, J., Olsen, H.E.: MinION analysis and reference consortium: Phase 1 data release and analysis. F1000 Research 4 (2015). doi:10.12688/f1000research.7201.1

29. Rhoads, A., Au, K.F.: Pacbio sequencing and its applications. Genomics, Proteomics and Bioinformatics 13(5), 278–289 (2015)

30. Jain, M., Olsen, H.E., Paten, B., Akeson, M.: The oxford nanopore minion: delivery of nanopore sequencing to the genomics community. Genome Biology 17(1), 239 (2016)

31. Jain, M., Fiddes, I.T., Miga, K.H., Olsen, H.E., Paten, B., Akeson, M.: Improved data analysis for the minion nanopore sequencer. Nature methods 12, 351–356 (2015)

32. Cretu Stancu, M., van Roosmalen, M.J., Renkens, I., Nieboer, M.M., Middelkamp, S., de Ligt, J., Pregno, G., Giachino, D., Mandrile, G., Espejo Valle-Inclan, J., Korzelius, J., de Bruijn, E., Cuppen, E., Talkowski, M.E., Marschall, T., de Ridder, J., Kloosterman, W.P.: Mapping and phasing of structural variation in patient genomes using nanopore sequencing. Nature Communications 8(1326) (2017). doi:10.1038/s41467-017-01343-4

33. Patterson, M., Marschall, T., Pisanti, N., van Iersel, L., Stougie, L., Klau, G.W., Schönhuth, A.: WhatsHap: Haplotype assembly for future-generation sequencing reads. In: RECOMB. LNCS, vol. 8394, pp. 237–249 (2014)

34. Patterson, M., Marschall, T., Pisanti, N., van Iersel, L., Stougie, L., Klau, G.W., Schönhuth, A.: WhatsHap: Weighted haplotype assembly for future-generation sequencing reads. Journal of Computational Biology 6(1), 498–509 (2015)

35. Pirola, Y., Zaccaria, S., Dondi, R., Klau, G.W., Pisanti, N., Bonizzoni, P.: HapCol: accurate and memory-efficient haplotype assembly from long reads. Bioinformatics 32(11), 1610–1617 (2016). doi:10.1093/bioinformatics/btv495

36. Bracciali, A., Aldinucci, M., Patterson, M., Marschall, T., Pisanti, N., Merelli, I., Torquati, M.: PWHATSHAP: efficient haplotyping for future generation sequencing. BMC Bioinformatics 17(11), 342 (2016). doi:10.1186/s12859-016-1170-y

37. Martin, M., Patterson, M., Garg, S., Fischer, S.O., Pisanti, N., Klau, G.W., Schoenhuth, A., Marschall, T.: WhatsHap: fast and accurate read-based phasing (2016)

38. Bonizzoni, P., Della Vedova, G., Dondi, R., Li, J.: The haplotyping problem: An overview of computational models and solutions. Journal of Computer Science and Technolgy 18(6), 675–688 (2003)

39. Kuleshov, V.: Probabilistic single-individual haplotyping. Bioinformatics 30(17), 379–385 (2014)

40. Zook, J.M.: Integrating human sequence data sets provides a resource of benchmark SNP and indel genotype calls. Nature Biotechnology 32, 246–251 (2014)

41. Zook, J.M., Catoe, D., McDaniel, J., Vang, L., Spies, N., Sidow, A., Weng, Z., Liu, Y., Mason, C.E., Alexander, N., Henaff, E., McIntyre, A.B.R., Chandramohan, D., Chen, F., Jaeger, E., Moshrefi, A., Pham, K., Stedman, W., Liang, T., Saghbini, M., Dzakula, Z., Hastie, A., Cao, H., Deikus, G., Schadt, E., Sebra, R., Bashir, A., Truty, R.M., Chang, C.C., Gulbahce, N., Zhao, K., Ghosh, S., Hyland, F., Fu, Y., Chaisson, M., Xiao, C., Trow, J., Sherry, S.T., Zaranek, A.W., Ball, M., Bobe, J., Estep, P., Church, G.M., Marks, P., Kyriazopoulou-Panagiotopoulou, S., Zheng, G.X.Y., Schnall-Levin, M., Ordonez, H.S., Mudivarti, P.A., Giorda, K., Sheng, Y., Rypdal, K.B., Salit, M.: Extensive sequencing of seven human genomes to characterize benchmark reference materials. Scientific Data 3(160025) (2016). doi:10.1038/sdata.2016.25

42. Kalman, L., Datta, V., Williams, M., Zook, J.M., Salit, M.L., Han, J.-Y.: Development and characterization of reference materials for genetic testing: Focus on public partnerships. Annals of Laboratory Medicine 36(6), 513–520 (2016). doi:10.3343/alm.2016.36.6.513

43. Garg, S., Martin, M., Marschall, T.: Read-based phasing of related individuals. Bioinformatics 32(12), 234–242 (2016)

44. Delaneau, O.: Haplotype estimation using sequencing reads. American Journal of Human Genetics 93, 687–696 (2013)

45. Hamada, Y.O.K.A.M.: PBSIM: Pacbio reads simluator–toward accurate genome assembly. Bioinformatics 29, 119–121 (2012)

46. Li, H.: Aligning sequence reads, clone sequences and assembly contigs with BWA-MEM. ArXiv e-prints (2013). 1303.3997

47. Fischer, S.O., Marschall, T.: Selecting Reads for Haplotype Assembly. bioRxiv, 046771 (2016). doi:10.1101/046771

48. Marschall, T. personal communication (2018)

49. Lo, C., Bashir, A., Bansal, V., Bafna, V.: Strobe sequence design for haplotype assembly. BMC Bioinformatics 12(Suppl. 1), 24 (2011)

50. Duitama, J., et al.: Fosmid-based whole genome haplotyping of a HapMap trio child: evaluation of single individual haplotyping techniques. Nucleic Acids Research 40, 2041–2053 (2012)

51. Hi-c: a comprehensive technique to capture the conformation of genomes. Methods 58(3), 268–276 (2012). doi:10.1016/j.ymeth.2012.05.001

